# High-Accuracy Protein Structures by Combining Machine-Learning with Physics-Based Refinement

**DOI:** 10.1101/731521

**Authors:** Lim Heo, Michael Feig

## Abstract

Protein structure prediction has long been available as an alternative to experimental structure determination, especially via homology modeling based on templates from related sequences. Recently, models based on distance restraints from co-evoluttionary analysis via machine learning have significantly expanded the ability to predict structures for sequences without templates. One such method, AlphaFold, also performs well on sequences were templates are available but without using such information directly. Here we show that combining machine-learning based models from AlphaFold with state-of-the-art physics-based refinement via molecular dynamics simulations further improves predictions to outperform any other prediction method tested during the latest round of CASP. The resulting models have highly accurate global and local structure, including high accuracy at functionally important interface residues, and they are highly suitable as initial models for crystal structure determination via molecular replacement.

## INTRODUCTION

High-resolution protein structures are key for understanding mechanistic details of biological function and for guiding rational drug design. The main experimental source of such structures is X-ray crystallography, nuclear magnetic resonance spectroscopy, or cryo-electronmicroscopy. By now, there is fairly comprehensive coverage of the manifold of protein structures encountered in biology, but a complete set of structures for any specific organism other than human immunodeficiency virus remains a very distant goal due to experimental limitations (Kolodny, Pereyaslavets, Samson, & Levitt, 2013; Zhang, Hubner, Arakaki, Shakhnovich, & Skolnick, 2006).

The computational modeling of protein structures has long been an alternative (Baker & Sali, 2001; Zhang, 2009). Structures can be predicted with reasonable accuracy by using templates based on sequence homology (Kryshtafovych et al., 2018). More recently, high-accuracy models have also been derived based on intramolecular distance restraints according to inferred co-evolutionary relationships (Kim, Dimaio, Yu-Ruei Wang, Song, & Baker, 2014; Ovchinnikov et al., 2017; Schaarschmidt, Monastyrskyy, Kryshtafovych, & Bonvin, 2018), especially for sequences where structural templates cannot be identified. Co-evolution analysis is highly suitable for the application of deep learning methods that are trained to predict residue interactions from multiple sequence alignments (Adhikari, Hou, & Cheng, 2018; Buchan & Jones, 2018; Hou, Wu, Cao, & Cheng, 2019; Jones & Kandathil, 2018; Kandathil, Greener, & Jones, 2019; Wang, Sun, Li, Zhang, & Xu, 2017; Wang, Sun, & Xu, 2018; Xu & Wang, 2019). This has been demonstrated most convincingly by DeepMind’s AlphaFold program during the recent 13^th^ Critical Assessment of Techniques for Protein Structure Prediction (CASP13) (AlQuraishi, 2019). AlphaFold outperformed all other methods for ‘free-modeling’ (FM) targets where templates were not available but was also competitive in the ‘template-based modeling’ (TBM) categories, although, presumably, without using templates directly (Abriata, Tamo, & Dal Peraro, 2019).

Predicted protein structures often capture secondary structures and fold topology correctly, but retain modeling errors compared to experimental structures (Feig, 2017). Reaching experimental accuracy is the goal of structure refinement techniques. Physics-based methods based on molecular dynamics (MD) simulations have been most successful in improving both global and local structure (Feig, 2016; Feig & Mirjalili, 2016; Heo & Feig, 2018a, 2018b; Mirjalili & Feig, 2013; Mirjalili, Noyes, & Feig, 2014; Read, Sammito, Kryshtafovych, & Croll, 2019). Current methods can provide consistent refinement for most structures while it is becoming increasingly possible to approach experimental accuracy based on root mean square deviations (RMSDs) of 1 Å or better as demonstrated during CASP13 (Heo, Arbour, & Feig, 2019). Interestingly, refinement was especially successful for two targets that came initially from AlphaFold predictions. This suggested that MD-based refinement could allow significant imrprovements of machine-learning based models due to complementarity between the data-driven and physics-based approaches. Indeed, we find that the application of MD-based structure refinement to AlphaFold’s machine learning models submitted during CASP13 leads to models that overall surpass the accuracy of other approaches, remarkably even for targets where traditional template-based modeling is possible. The combination of machine learning and physics-based refinement therefore results in the highest-resolution protein structure models to date.

## RESULTS AND DISCUSSION

We applied the physics-based model refinement protocol established during CASP13 to “model 1” predictions from AlphaFold (group A7D during CASP13) for all targets. The refinement protocol is based on MD simulations and does not apply any knowledge of the native structure or other data (*see Methods*).

### Model accuracy

The model qualities for the generated models were evaluated according to the CASP13 assessment procedures, i.e. TBM-score and FM-score for targets in the TBM and FM categories, respectively *(see Methods)*. Using the same scores as during CASP13 allowed us to directly compare with results from other groups without having to reevaluate all of the predictions. **Figure 1** summarizes the scores for the AlphaFold models before and after refinement in comparison with models from other groups generated with different methods. As reported during CASP13, AlphaFold models were very competitive, ranking first in the FM category and 4^th^ in the TBM category. MD refinement further improved AlphaFold predictions. TBM targets were refined most, resulting in an increased TBM-score of 61.7 (from 44.6) that surpassed the performance of the best template-based modeling protocols. The improvements after refinement were greater for TBM-easy targets (36.0 vs. 24.2) than for TBM-hard targets (25.6 vs. 20.3), where ‘easy’ and ‘hard’ relate to the availability and quality of suitable templates. For FM targets, where templates are not available, refinement improved AlphaFold models moderately with an increased FM-score from 67.2 to 69.0. Improvements due to refinement were consistent across different targets. 78 out of 104 (75%) of targets were refined, including 34 out of 40 (85%) TBM-easy targets.

**Figure 1.**
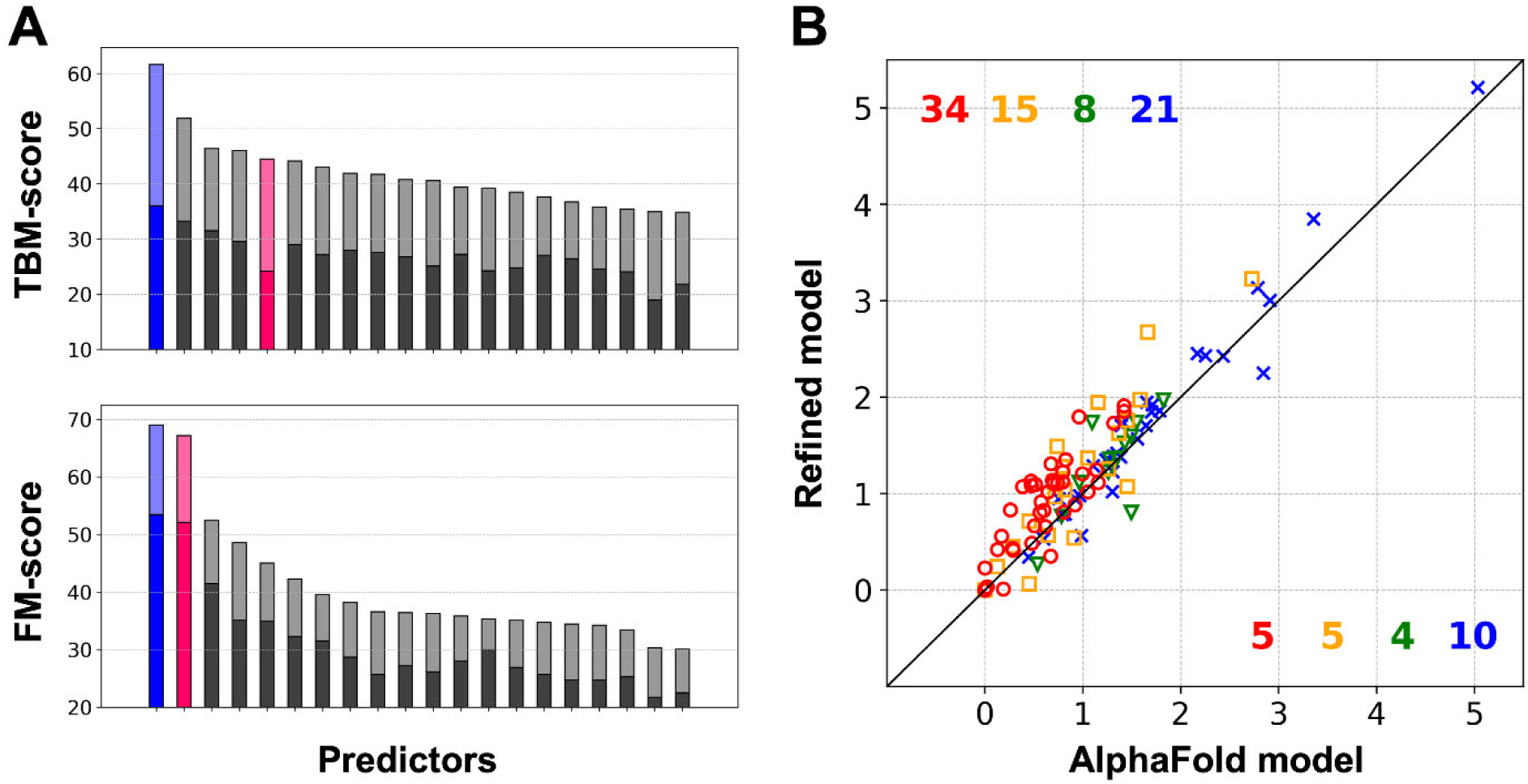
Assessment of refined AlphaFold models in comparison with CASP13 results. (A) Scores (sum of assessor’s formula) for TBM (top) and FM (bottom) targets. Results for AlphaFold (A7D) before and after refinement are shown in red and blue, respectively. Other top-performing groups are shown in grey. TBM-easy and FM-hard targets are shown in darker colors, while TBM-hard and FM/TBM targets are shown in lighter colors. (B) Head-to-head comparison between AlphaFold models before and after refinement based on assessor’s formula. Each point corresponds to a target. Targets for TBM-easy, TBM-hard, FM/TBM, and FM-hard are depicted in red circles, orange squares, green triangles, and blue Xs, respectively. The number of better targets in each subcategory is shown at the top (refined model is better) and bottom (AlphaFold model is better) in the same colors.

Changes in individual quality measures after refinement (**Figure 1 – Supplementary 1** and **Table S1)** indicate that improvements were not limited to specific structure features, but that there was an overall enhancement in global and local structural quality as well as better residue-wise error estimation. During CASP13, AlphaFold generated the best overall models of any group for 25 out of 104 (24%) targets. AlphaFold+refinement resulted in the best models for 41 targets (39%) (*see* **Figure 1 – Supplementary 2**). Interestingly, even though AlphaFold or our refinement method did not use templates, it was possible to generate the best models for 14 TBM-easy and 8 TBM-hard targets, respectively, significantly improving upon the accuracy of the unrefined AlphaFold models and surpassing the average performance of any other group in all categories based on Z-scores (*see* **Figure 1 – Supplementary 3–4, Table S2 and S3**). Only in the TBM-easy category and only with respect to GDT-HA scores were the refined AlphaFold models exceeded by models from the Zhang group.

AlphaFold’s deep-learning method focuses primarily on predicted inter-residue distance distributions and backbone dihedral angles and the accuracy in the resulting models is essentially limited to the residue level. Moreover, model errors likely reflect structural variations within in a given protein family since contacts are obtained from deep multiple sequence alignments. This leaves room for physics-based approaches to provide significant refinement at the atomistic level. As may be expected, the accuracy of AlphaFold models depends on how many homologous sequences were available (see **Figure 1 – Supplementary 5A**) while MD-based refinement success is independent of available homologous sequences (**Figure 1 – Supplementary 5B**), further highlighting the complementarity of both approaches.

**Figure 2** illustrates two typical examples of successful refinement: The initial AlphaFold models have correct folds, but they slightly mis-predict some regions such as loops and display suboptimal packing between side-chains. These issues were resolved by the refinement. Loop structures are presumably more difficult to predict by AlphaFold because there are fewer inter-residue contacts compared to secondary structure elements. On the other hand, the exact packing of side chains due to physical laws may be difficult to obtain simply based on sequence-derived contacts.

**Figure 2.**
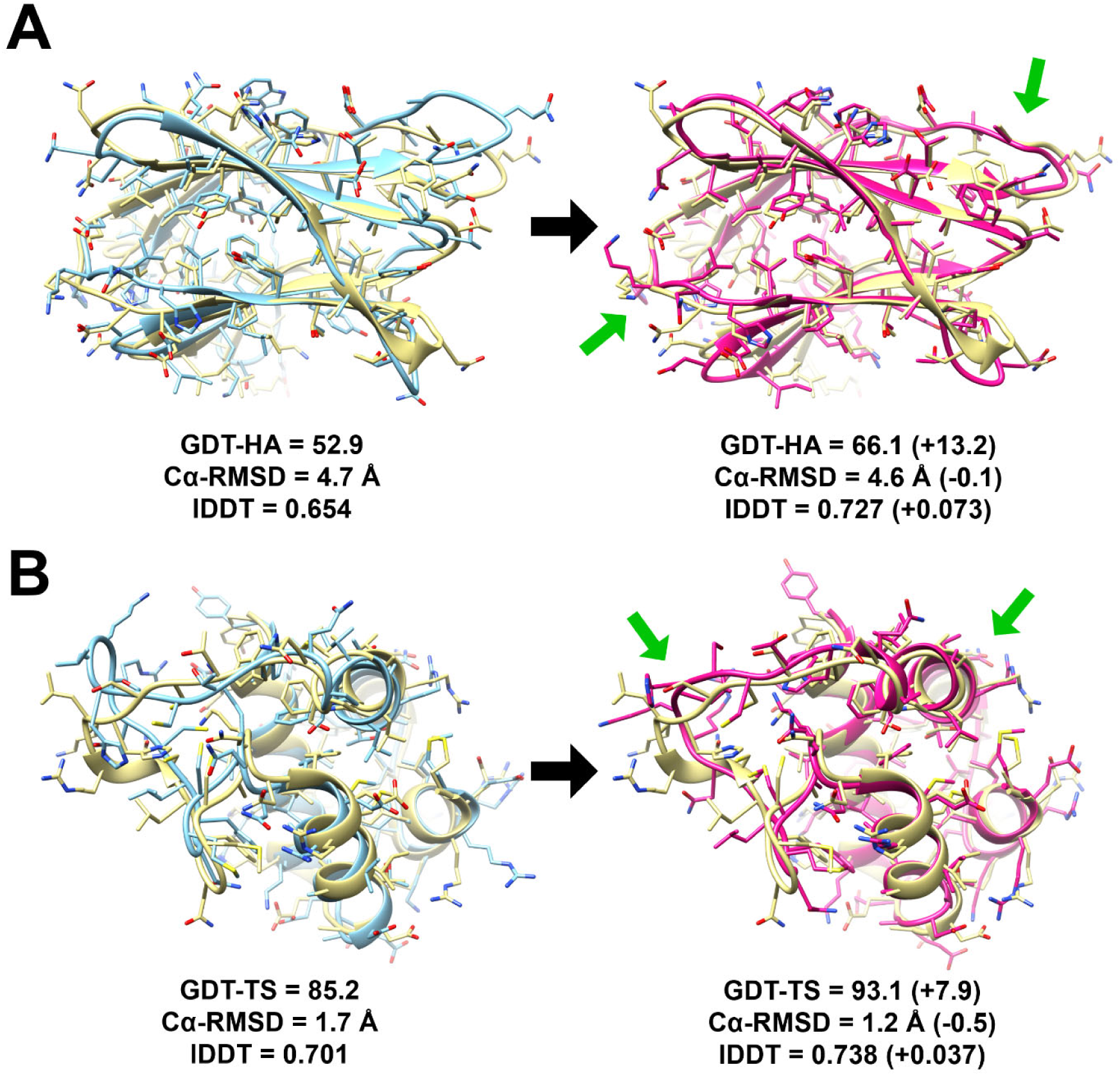
Examples of successful refined models with (A) T0964-D1 for TBM category and (B) T0990-D1 for FM category. AlphaFold models and their refined models are shown in blue and magenta, respectively. Experimental structures are overlayed in yellow after superposition. Regions that were most significantly improved after refinement are indicated by green arrows. Model quality metrics are shown below each model.

### Accuracy of protein-protein interfaces

Protein-protein interactions are important in the function of many proteins (Negri et al., 2010; Pawson & Nash, 2000; Russell et al., 2004). There were 27 hetero- and 57 homo-oligomeric targets allowing the separate evaluation of the accuracy of interface residues (defined as residues with a heavy atom distance closer than 10 Å to another protein). The interface-RMSD (iRMSD) based on the backbone atoms of interface residues was lower than 4 Å in either initial or refined AlphaFold models for 14 hetero- and 22 homo-oligomeric targets. Among these targets, refinement improved not just the global structure (in terms of GDT-HA), but also the interface (in terms of iRMSD) (**Figure 3A**). On average, iRMSD values were decreased by 0.15 Å (p=4.1%, n=36) with improvements for 65% of the targets. **Figure 3B** illustrates a successful example of a signficanctly refined interface even although structures were refined as monomers. Two targets (T1015s1-D1 and T1019s1-D1) became significantly worse after refinement in terms of GDT-HA and iRMSD. These targets have a typical Zinc finger moiety but a Zinc ion was not included in the refinement. If these targets are excluded, the improvement in iRMSD increases to 0.22 Å (p=0.14%, n=34).

**Figure 3.**
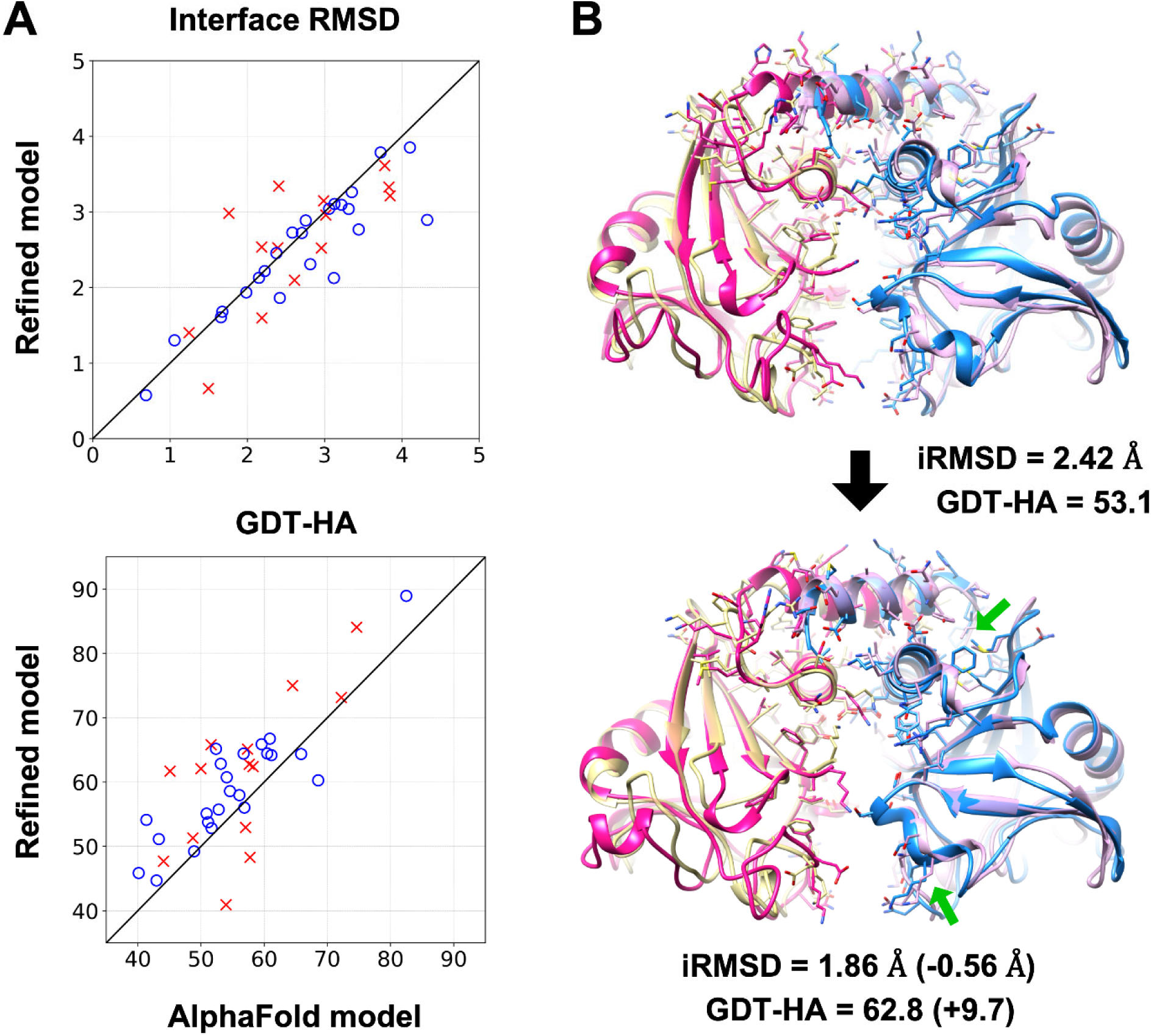
Refinement at protein-protein interfaces. (A) Structure quality comparisons between AlphaFold and refined models in terms of interface RMSD (top) and GDT-HA (bottom). Hetero-and homo-olligomers are shown as red Xs and blue circles, respectively. (B) An example of refinement at the interface of T0997-D1. Both AlphaFold (top) and refined models (bottom) are shown in red and blue, while the experimental structure is shown in yellow and pink. Structures are depicted in cartoon representations, and interface residues are also shown as sticks. Models were superimposed onto the experimental structures by aligning the interface residues. Significantly improved interface regions are indicated by green arrows. Model qualities measures are given below each model.

### Use of models for molecular replacement

A practical value of predicted structures is their use as starting models for solving crystal structures via molecular replacement (MR).(Scapin, 2013) We followed the procedure that was used in the CASP13 assessment where MR success in Phaser(McCoy et al., 2007) was evaluated in terms of log-liklihood-gains (LLG). There are 19 crystallographic data sets available so far for a total of 27 CASP13 targets. LLG scores increased for most targets after refinement (**Table S2**). LLG scores of 120 or higher often indicate the possibility for successful MR.(McCoy et al., 2007) Between initial and refined AlphaFold models, this was reached for 14 targets and out of those the refined models had higher LLG than the AlphaFold model for 10 targets making success with MR more likely. Typically, MR relies on homology models with high sequence identity templates or MR-Rosetta(DiMaio, 2013) for sequences with lower sequence identity templates. The results presented here suggest that refined machine learning-based models are becoming a good alternative for solving crystallographic data via MR.

## CONCLUSIONS

Co-evolutionary analysis via machine learning has significantly improved prediction accuracies for sequences where templates are not available but has also been demonstrated to perform well for sequences where templates could have been used instead. Here we show that physics-based refinement can further improve the accuracy of machine-learning models to exceed the accuracy of any other available method based on the targets that were assessed in CASP13. This suggests that a protocol that combines machine-learning to obtain residue-residue distance information and torsional preferences with distance-geometry model generation (such as in AlphaFold) followed by MD-based refinement to improve structural details at the atomistic level is emerging as a universal protocol for obtaining the most accurate predictions irrespective of whether template structures are available for a specific sequence or not. Additional analysis shows that functionally important residues at protein-protein interfaces can be predicted at high accuracy and that refined AlphaFold models are highly useful as search models for molecular replacement during crystallographic structure determination.

## METHODS

### Refinement of AlphaFold Models

The latest version of the PREFMD protocol (Heo et al., 2019) was applied to refine AlphaFold (A7D)’s “model 1”s generated during CASP13 for protein tertiary structure predictions. The AlphaFold models were downloaded from the CASP web page (http://predictioncenter.org/download_area/CASP13/). The overall refinement method consisted of pre-sampling, sampling, and post-sampling stages. Details are described elsewhere (Heo et al., 2019) and only the key points are briefly outlined here. In the pre-sampling stage, local streochemical errors such as atomic clashes and poor backbone dihedral angles were resolved by locPREFMD (Feig, 2016). At the sampling stage, molecular dynamics (MD) simulations were conducted to explore conformational space. One of two variants was chosen based on target size. In the full iterative protocol, three iterations of MD simulations were carried out, while a more conservative protocol consisted of only a single iteration. For every iteration in the iterative protocol, MD simulations were started from new initial structures chosen from the previous iteration. In the iterative protocol, flat-bottom harmonic restraints were used to allow significant structure changes from the initial model within a certain raidus. This protocol was used for smaller targets. In the conservative protocol, a harmonic restraint function was used to achieve more consistent, but moderate refinement for larger targets where extensive sampling would have been a challenge. The latest version of CHARMM force field, c36m (Huang et al., 2017), and its in-house modification (Heo et al., 2019) were used for both sampling protocols. After the sampling stage, generated snapshots were evaluated by using the Rosetta score (Park et al., 2016). Conformations with low Rosetta scores were then subjected to ensemble averaging and locPREFMD was applied again to the ensemble averaged structures. Finally, residuewise errors were estimated by using root-mean-square-fluctuation (RMSF) from short MD simulations. We note that in the original method used during CASP13 we attempted to identify putative ligands, but this was not done here in order to implement a fully automatic protocol.

### CASP13 assessment scores

For template-based modeling (TBM) category targets, the weighted Z-score sum of Global Distance Test-High Accuracy (GDT-HA) (Zemla, 2003), Local Distance Difference Test (lDDT) (Mariani, Biasini, Barbato, & Schwede, 2013), Contact Area Difference-all atom (CAD-aa) (Olechnovic, Kulberkyte, & Venclovas, 2013), SphereGrinder (Kryshtafovych, Monastyrskyy, & Fidelis, 2014), and Accuracy Self Estimate (ASE) scores (Kryshtafovych et al., 2016) were used (Kryshtafovych et al., 2018). This score considers global structure similarity (GDT-HA), local structure similarity (lDDT, CAD-aa, and SphereGrinder), as well as estimates of residue-wise model errors. Free modeling (FM) category targets were assessed separately in CASP13 using a different score that consists of the weighted Z-score sum of Global Distance Test-Tertiary Structure (GDT-TS) (Zemla, 2003) and Quality Control Score (QCS) (Cong et al., 2011), both of which focus only on global structure similarity (Abriata et al., 2019).

### Homologous Sequence Search

To investigate correlations between the number of homologous sequence for a target and the quality of models from AlphaFold and its refined model we searched for homlogous sequences based on the Uniclust30 database (clustered sequences of UniProtKB at 30% pairwise sequence identity) (Mirdita et al., 2017). HHblits (Remmert, Biegert, Hauser, & Soding, 2011) was used for the sequence search with default options except for the number of iterations set to 3. Sequences with a very higher pairwise sequence identity (above 90%) were filtered out using *hhfilter*, a program from HHsuite (Soding, 2005; Steinegger et al., 2019). Results were compared based on he Uniclust30 databases released in August 2018 and September 2016, but no significant differences were observed. The results reported here are based on the August 2018 release.

### Protein Models as Molecular Replacement Search Models

Crystallographic data sets were downloaded from the Protein Data Bank (PDB) (Westbrook, Feng, Chen, Yang, & Berman, 2003) and prepared by using Phenix (Adams et al., 2010). Molecular replacement (MR) was conducted with Phaser (McCoy et al., 2007) to evaluate log-liklihood gains (LLG) of protein models. We did not perform full MR searches to reduce the computational costs with the large number of models that were considered. Instead, we carried out rigid-body refinement after superimposition of a model on its corresponding component in the crystal structure. If there were multiple different components in the crystal, the model was evaluated after the other components were placed as background structures, and the LLG were calculated with the difference made by the model. Predicted residuewise model errors *e* were utilized as B-factors calculated as 8π^2^/3 *e*^2^. We report the best LLG between the MR results with and without the predicted errors for a given model.

## ACKNOWLEDGEMENTS

We are grateful to the CASP organizers and assessors for organizing CASP13 and we acknowledge DeepMind for the valuable AlphaFold model predictions. We thank Prof. Read for providing scripts for molecular replacement with Phaser. This research was supported by National Institutes of Health Grants R01 GM084953 and R35 GM126948. Computational resources were used at the National Science Foundation’s Extreme Science and Engineer-ing Discovery Environment (XSEDE) facilities under Grant TG-MCB090003.

