## Supplemental Tables and Figures for "High-Accuracy Protein Structures by Combining Machine-Learning with Physics-Based Refinement"

**Table S1. Comparison between AlphaFold and its refined models**

| Quality Measure | Category | Number of Targets | AlphaFold[1] | Refined[1] | ΔMeasure | P-value[1] |
|---|---|---|---|---|---|---|
| Z-score | Overall | 104 | 1.073 | **1.249** | 0.176 | $1.2\times10^{-8}$ |
| | TBM-easy | 40 | 0.605 | **0.876** | 0.271 | $6.3\times10^{-8}$ |
| | TBM-hard | 21 | 0.969 | **1.209** | 0.240 | $3.8\times10^{-3}$ |
| | FM/TBM | 12 | 1.255 | 1.290 | 0.035 | 0.35 |
| | FM | 31 | 1.676 | 1.741 | 0.066 | $5.4\times10^{-2}$ |
| GDT-HA | Overall | 104 | 46.70 | **49.35** | 2.65 | $3.0\times10^{-7}$ |
| | TBM-easy | 40 | 55.32 | **58.31** | 2.99 | $2.2\times10^{-4}$ |
| | TBM-hard | 21 | 44.61 | **47.76** | 3.14 | $3.7\times10^{-3}$ |
| | FM/TBM | 12 | 49.60 | 52.25 | 2.65 | $9.9\times10^{-2}$ |
| | FM | 31 | 35.87 | **37.76** | 1.89 | $2.0\times10^{-2}$ |
| lDDT | Overall | 104 | 0.6022 | **0.6143** | 0.0121 | $2.0\times10^{-5}$ |
| | TBM-easy | 40 | 0.6665 | **0.6848** | 0.0183 | $4.6\times10^{-4}$ |
| | TBM-hard | 21 | 0.6238 | **0.6387** | 0.0150 | $2.6\times10^{-3}$ |
| | FM/TBM | 12 | 0.5940 | 0.5916 | -0.0024 | 0.41 |
| | FM | 31 | 0.5077 | **0.5156** | 0.0079 | $2.9\times10^{-2}$ |
| CAD-aa | Overall | 104 | 0.6234 | **0.6365** | 0.0131 | $4.0\times10^{-9}$ |
| | TBM-easy | 40 | 0.6608 | **0.6794** | 0.0186 | $7.9\times10^{-7}$ |
| | TBM-hard | 21 | 0.6326 | **0.6493** | 0.0167 | $5.8\times10^{-4}$ |
| | FM/TBM | 12 | 0.6076 | 0.6104 | 0.0029 | 0.36 |
| | FM | 31 | 0.5751 | **0.5826** | 0.0075 | $1.3\times10^{-2}$ |
| SphGr | Overall | 104 | 72.23 | 72.44 | 0.21 | 0.23 |
| | TBM-easy | 40 | 81.77 | 82.36 | 0.59 | 0.099 |
| | TBM-hard | 21 | 75.09 | 74.77 | -0.32 | 0.28 |
| | FM/TBM | 12 | 71.79 | 70.92 | -0.87 | 0.27 |
| | FM | 31 | 58.16 | 58.66 | 0.49 | 0.11 |
| ASE | Overall | 104 | 65.86 | **83.89** | 18.03 | $4.5\times10^{-10}$ |
| | TBM-easy | 40 | 59.16 | **88.84** | 29.68 | $1.1\times10^{-7}$ |
| | TBM-hard | 21 | 63.08 | **82.12** | 19.04 | $1.3\times10^{-3}$ |
| | FM/TBM | 12 | 82.19 | 85.92 | 3.73 | 0.15 |
| | FM | 31 | 70.05 | **77.91** | 7.86 | 0.033 |
| Cα-RMSD | Overall | 104 | 6.652 | **6.536** | -0.116 | $2.4\times10^{-4}$ |
| | TBM-easy | 40 | 4.431 | **4.288** | -0.144 | $2.3\times10^{-3}$ |
| | TBM-hard | 21 | 8.784 | 8.713 | -0.071 | 0.11 |
| | FM/TBM | 12 | 3.910 | 3.809 | -0.102 | 0.21 |
| | FM | 31 | 9.135 | **9.019** | -0.116 | 0.047 |
| GDT-TS | Overall | 104 | 64.42 | **66.28** | 1.86 | $3.8\times10^{-6}$ |
| | TBM-easy | 40 | 73.44 | **75.35** | 1.91 | $1.4\times10^{-3}$ |
| | TBM-hard | 21 | 62.08 | **64.43** | 2.35 | $4.1\times10^{-3}$ |
| | FM/TBM | 12 | 68.57 | 70.51 | 1.94 | 0.15 |
| | FM | 31 | 52.74 | **54.19** | 1.45 | 0.017 |
| QCS | Overall | 104 | 81.80 | 82.08 | 0.28 | 0.12 |
| | TBM-easy | 40 | 88.98 | 89.44 | 0.47 | 0.093 |
| | TBM-hard | 21 | 80.33 | 80.11 | -0.23 | 0.36 |
| | FM/TBM | 12 | 80.64 | 80.32 | -0.32 | 0.39 |
| | FM | 31 | 73.98 | **74.60** | 0.62 | 0.030 |

[1] Significantly improved (P-value < 0.05) quality measures are highlighted in blue. P-values were obtained from Student's paired T-test (one-tailed). Note that none of the quality measures became significantly worse.

**Table S2. Comparison between AlphaFold models and other top prediction groups**

| Quality Measures | Category | Number of targets | AlphaFold | Best[1] | Zhang[1] | Seok-refine[1] | Zhang-server[1] | QUARK[1] |
|---|---|---|---|---|---|---|---|---|
| Z-score | Overall | 104 | 1.073 | 1.543 | 1.004 | 0.726 | 0.849 | 0.857 |
| | TBM-easy | 40 | 0.605 | 1.079 | 0.832 | 0.788 | 0.740 | 0.727 |
| | TBM-hard | 21 | 0.969 | 1.392 | 0.889 | 0.711 | 0.786 | 0.718 |
| | FM/TBM | 12 | 1.255 | 1.554 | 0.917 | 0.683 | 0.833 | 0.841 |
| | FM | 31 | 1.676 | 2.239 | 1.335 | 0.672 | 1.039 | 1.126 |
| GDT-HA | Overall | 104 | 46.70 | 54.24 | 46.24 | 42.36 | 43.91 | 43.80 |
| | TBM-easy | 40 | 55.32 | 65.96 | 61.31 | 57.55 | 58.81 | 58.81 |
| | TBM-hard | 21 | 44.61 | 52.28 | 44.42 | 40.94 | 42.98 | 42.24 |
| | FM/TBM | 12 | 49.60 | 55.24 | 42.55 | 40.51 | 40.86 | 41.27 |
| | FM | 31 | 35.87 | 40.07 | 29.46 | 24.46 | 26.50 | 26.47 |
| lDDT | Overall | 104 | 0.6022 | 0.6463 | 0.5771 | 0.5586 | 0.5613 | 0.5578 |
| | TBM-easy | 40 | 0.6665 | 0.7318 | 0.6735 | 0.6780 | 0.6572 | 0.6535 |
| | TBM-hard | 21 | 0.6238 | 0.6567 | 0.5881 | 0.5781 | 0.5776 | 0.5710 |
| | FM/TBM | 12 | 0.5940 | 0.6175 | 0.5225 | 0.5142 | 0.5075 | 0.5117 |
| | FM | 31 | 0.5077 | 0.5400 | 0.4665 | 0.4084 | 0.4474 | 0.4432 |
| CAD-aa | Overall | 104 | 0.6234 | 0.6514 | 0.5861 | 0.5978 | 0.5747 | 0.5725 |
| | TBM-easy | 40 | 0.6608 | 0.7090 | 0.6508 | 0.6745 | 0.6348 | 0.6358 |
| | TBM-hard | 21 | 0.6326 | 0.6538 | 0.5876 | 0.6052 | 0.5781 | 0.5748 |
| | FM/TBM | 12 | 0.6076 | 0.6275 | 0.5442 | 0.5667 | 0.5442 | 0.5358 |
| | FM | 31 | 0.5751 | 0.5848 | 0.5177 | 0.5058 | 0.5068 | 0.5035 |
| SphGr | Overall | 104 | 72.23 | 78.22 | 69.17 | 62.60 | 66.48 | 66.04 |
| | TBM-easy | 40 | 81.77 | 89.96 | 84.59 | 80.74 | 81.70 | 81.14 |
| | TBM-hard | 21 | 75.09 | 79.16 | 70.17 | 64.72 | 68.62 | 67.21 |
| | FM/TBM | 12 | 71.79 | 74.45 | 60.35 | 59.11 | 58.71 | 57.20 |
| | FM | 31 | 58.16 | 63.91 | 52.00 | 39.12 | 48.40 | 49.17 |

[1] Significantly better or worse values than the AlphaFold models in each quality measure are highlighted in blue and red, respectively. P-values were obtained from Student's paired T-test (one-tailed), and P-values of 0.05 or less are used as the criterion for significance.

**Table S3. Comparison between refined AlphaFold models and other top prediction groups**

| Quality Measures | Category | Number of targets | Refined | Best[1] | Zhang[1] | Seok-refine[1] | Zhang-server[1] | QUARK[1] |
|---|---|---|---|---|---|---|---|---|
| Z-score | Overall | 104 | 1.249 | 1.543 | 1.004 | 0.726 | 0.849 | 0.857 |
| | TBM-easy | 40 | 0.876 | 1.079 | 0.832 | 0.788 | 0.740 | 0.727 |
| | TBM-hard | 21 | 1.209 | 1.392 | 0.889 | 0.711 | 0.786 | 0.718 |
| | FM/TBM | 12 | 1.290 | 1.554 | 0.917 | 0.683 | 0.833 | 0.841 |
| | FM | 31 | 1.741 | 2.239 | 1.335 | 0.672 | 1.039 | 1.126 |
| GDT-HA | Overall | 104 | 49.35 | 54.24 | 46.24 | 42.36 | 43.91 | 43.80 |
| | TBM-easy | 40 | 58.31 | 65.96 | 61.31 | 57.55 | 58.81 | 58.81 |
| | TBM-hard | 21 | 47.76 | 52.28 | 44.42 | 40.94 | 42.98 | 42.24 |
| | FM/TBM | 12 | 52.25 | 55.24 | 42.55 | 40.51 | 40.86 | 41.27 |
| | FM | 31 | 37.76 | 40.07 | 29.46 | 24.46 | 26.50 | 26.47 |
| lDDT | Overall | 104 | 0.6143 | 0.6463 | 0.5771 | 0.5586 | 0.5613 | 0.5578 |
| | TBM-easy | 40 | 0.6848 | 0.7318 | 0.6735 | 0.6780 | 0.6572 | 0.6535 |
| | TBM-hard | 21 | 0.6387 | 0.6567 | 0.5881 | 0.5781 | 0.5776 | 0.5710 |
| | FM/TBM | 12 | 0.5916 | 0.6175 | 0.5225 | 0.5142 | 0.5075 | 0.5117 |
| | FM | 31 | 0.5156 | 0.5400 | 0.4665 | 0.4084 | 0.4474 | 0.4432 |
| CAD-aa | Overall | 104 | 0.6365 | 0.6514 | 0.5861 | 0.5978 | 0.5747 | 0.5725 |
| | TBM-easy | 40 | 0.6794 | 0.7090 | 0.6508 | 0.6745 | 0.6348 | 0.6358 |
| | TBM-hard | 21 | 0.6493 | 0.6538 | 0.5876 | 0.6052 | 0.5781 | 0.5748 |
| | FM/TBM | 12 | 0.6104 | 0.6275 | 0.5442 | 0.5667 | 0.5442 | 0.5358 |
| | FM | 31 | 0.5826 | 0.5848 | 0.5177 | 0.5058 | 0.5068 | 0.5035 |
| SphGr | Overall | 104 | 72.44 | 78.22 | 69.17 | 62.60 | 66.48 | 66.04 |
| | TBM-easy | 40 | 82.36 | 89.96 | 84.59 | 80.74 | 81.70 | 81.14 |
| | TBM-hard | 21 | 74.77 | 79.16 | 70.17 | 64.72 | 68.62 | 67.21 |
| | FM/TBM | 12 | 70.92 | 74.45 | 60.35 | 59.11 | 58.71 | 57.20 |
| | FM | 31 | 58.66 | 63.91 | 52.00 | 39.12 | 48.40 | 49.17 |

[1] Significantly better or worse values than for the refined AlphaFold models in each quality measure are highlighted in blue and red, respectively. P-values were obtained from Student's paired T-test (one-tailed), and P-values of 0.05 or less were used as the criterion for significance.

**Table S4. Summary of molecular replacement results with protein models**

| PDB ID | Space Group | Target ID | # in ASU[1] | AlphaFold model 1 | | Best refined model[2] | | Best CASP13 model[3] | | |
|---|---|---|---|---|---|---|---|---|---|---|
| | | | | Cα-RMSD | LLG[4] | Cα-RMSD | LLG[4] | Cα-RMSD | LLG[4] | Model |
| 6cvz | $P\ 3_2$ | T0954-D1 | 3 | 2.8 | **182.8** | 2.6 | **425.7** | 2.8 | **285.0** | TS043_2 |
| 6btc | $P\ 6_1\ 2\ 2$ | T0958-D1 | 2 | 2.8 | 53.1 | 3.3 | 73.0 | 1.7 | **122.0** | TS431_1 |
| 6d2v | $P\ 2_1\ 2_1\ 2_1$ | T0965-D1 | 2 | 2.8 | **164.3** | 2.4 | **181.1** | 2.8 | **164.3** | TS043_1 |
| 5w6l | I 2 3 | T0966-D1 | 2 | 23.7 | 19.6 | 23.9 | 18.1 | 3.0 | **199.9** | TS117_1 |
| 6cci | $P\ 2_1\ 2_1\ 2_1$ | T0969-D1 | 1 | 6.1 | 31.2 | 6.1 | 27.8 | 6.7 | 36.9 | TS043_3 |
| 6g57 | $P\ 6_1\ 2\ 2$ | T0970-D1 | 4 | 3.8 | 45.5 | 3.7 | 41.9 | 3.2 | **292.6** | TS043_4 |
| 6hrh | C 1 2 1 | T1003-D1 | 2 | 13.9 | **171.2** | 13.9 | 53.2 | 6.1 | **1769.1** | TS214_5 |
| 6q64 | $P\ 3_2\ 2\ 1$ | T1005-D1 | 1 | 6.5 | 43.7 | 6.5 | 43.3 | 7.8 | 70.4 | TS426_5 |
| 6qek | P 61 | T1006-D1 | 2 | 0.8 | **214.6** | 0.8 | **464.0** | 0.6 | **662.2** | TS457_1 |
| 6dru | $P\ 3_2\ 2\ 1$ | T1009-D1 | 2 | 4.9 | **136.8** | 4.9 | **129.8** | 4.9 | **621.6** | TS214_1 |
| 6e4b | C 1 2 1 | T1016-D1 | 4 | 3.4 | **601.6** | 3.4 | **983.4** | 1.9 | **1184.5** | TS043_2 |
| 6n91 | $P\ 3_2\ 2\ 1$ | T1018-D1 | 2 | 2.0 | **259.3** | 2.0 | **367.6** | 1.8 | **1206.6** | TS086_5 |
| 6f45[5] | $C\ 2\ 2\ 2_1$ | T0953s1-D1 | 3 | 13.6 | 57.1 | 14.1 | 66.4 | 12.4 | 70.8 | TS221_1 |
| | | T0953s2-D1 | 1 | 5.9 | 19.7 | 5.9 | 16.9 | 6.5 | **160.9** | TS145_1 |
| | | T0953s2-D2 | 1 | 6.1 | **133.1** | 6.6 | 35.2 | 6.1 | **133.1** | TS043_1 |
| | | T0953s2-D3 | 1 | 9.5 | 15.8 | 10.2 | 10.1 | 10.7 | 22.8 | TS274_5 |
| 6cp8[5] | $P\ 1\ 2_1\ 1$ | T0957s1-D1 | 2 | 13.3 | 95.9 | 13.4 | 67.7 | 12.2 | **124.3** | TS043_2 |
| | | T0957s1-D2 | 2 | 1.9 | **173.5** | 1.9 | **242.3** | 1.7 | **264.5** | TS043_4 |
| | | T0957s2-D1 | 2 | 4.6 | 34.7 | 4.3 | 30.5 | 9.8 | 51.5 | TS112_4 |
| 6cp9[5] | $P\ 2_1\ 2_1\ 2$ | T0968s1-D1 | 4 | 4.4 | 64.4 | 4.3 | **142.1** | 4.3 | 85.5 | TS043_3 |
| | | T0968s2-D1 | 4 | 2.3 | **288.5** | 2.3 | **260.2** | 2.3 | **288.5** | TS043_1 |
| 6mxv[5] | $P\ 1\ 2_1\ 1$ | T0976-D1 | 2 | 1.6 | **533.5** | 1.5 | **763.3** | 1.5 | **1296.6** | TS068_1 |
| | | T0976-D2 | 2 | 2.8 | **977.6** | 2.7 | **1310.8** | 3.9 | **1386.5** | TS068_1 |
| 6gnx[5] | $P\ 3_2\ 2\ 1$ | T0980s1-D1 | 2 | 16.6 | 17.8 | 16.6 | 31.6 | 4.6 | 46.7 | TS043_3 |
| 6d7y[5] | $P\ 6_3$ | T0986s1-D1 | 1 | 2.6 | 4.9 | 2.1 | 24.3 | 6.1 | 28.5 | TS068_5 |
| | | T0986s2-D1 | 1 | 3.4 | 7.5 | 3.4 | 12.6 | 6.3 | 17.6 | TS498_1 |
| 6m9t[5] | C 1 2 1 | T1011-D1 | 1 | 3.8 | **373.8** | 3.8 | **456.2** | 3.5 | **450.1** | TS043_3 |

[1] Number of molecules in the asymmetric unit (ASU).

[2] The highest LLG score among the 5 refined models and Cα-RMSD for the corresponding model.

[3] The highest LLG score among the all CASP13 models and Cα-RMSD for the corresponding model.

[4] LLG scores above 120 are highlighted in bold characters. LLG score above 120 and better results before or after refinement are shaded in grey.

[5] Multiple different components in an asymmetric unit.

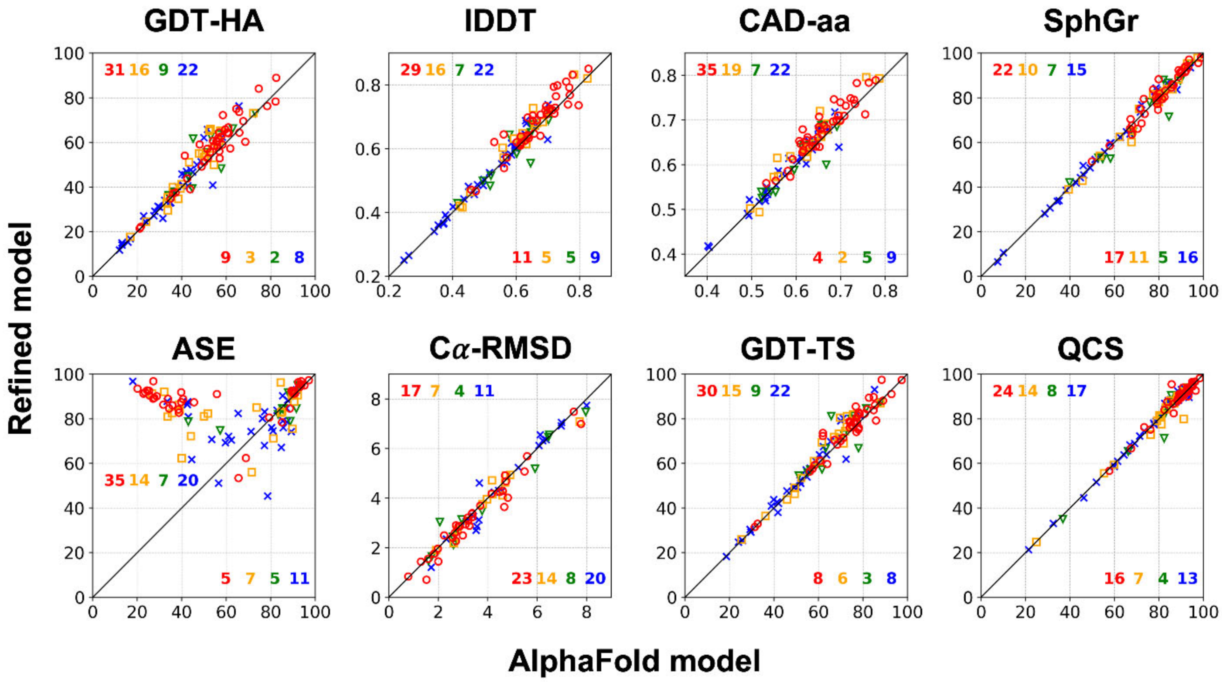


**Figure 1 – Supplementary 1.** Head-to-head comparison between initial and refined AlphaFold models for different quality measures. *See* **Figure 1B**.

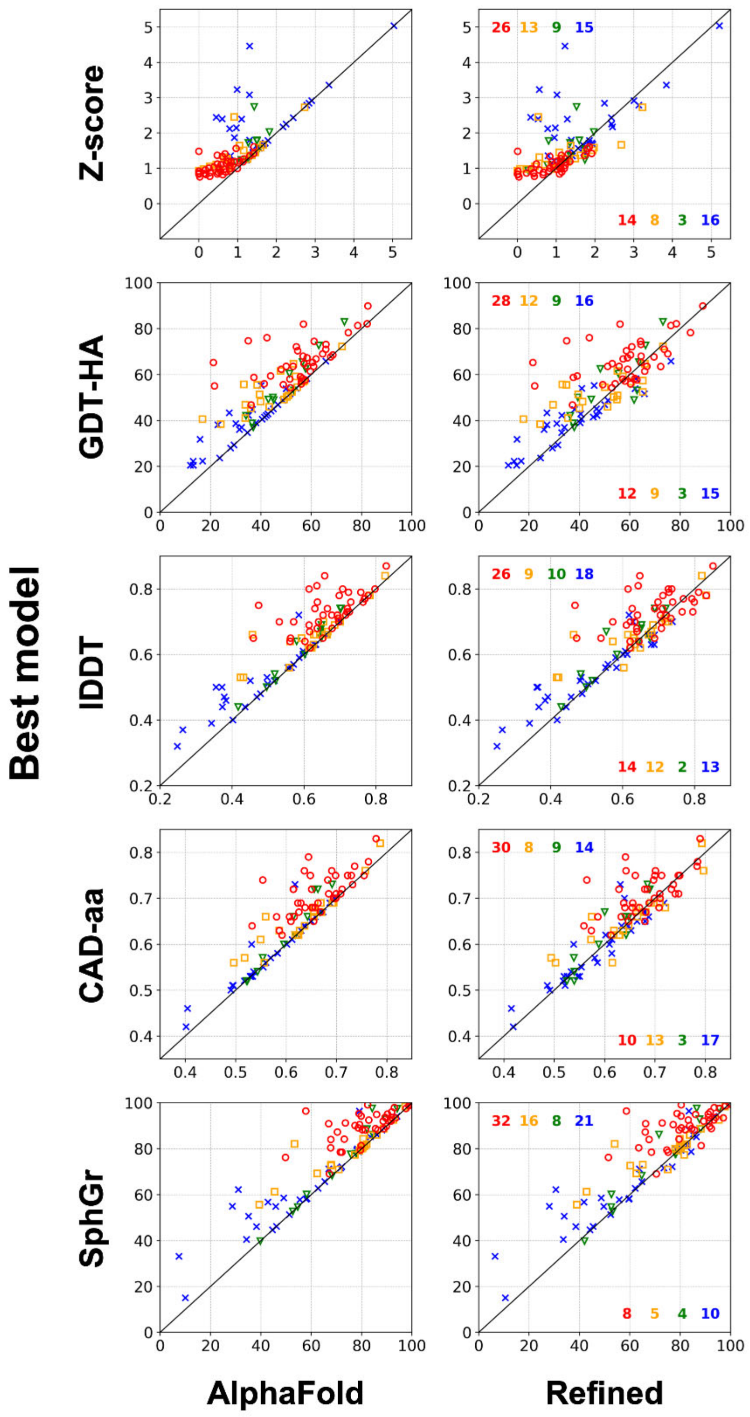


**Figure 1 – Supplementary 2.** Head-to-head comparison between initial and refined AlphaFold models and the best models for each target based on different quality measures. *See* **Figure 1B** and **Figure 1 - Supplementary 1**.

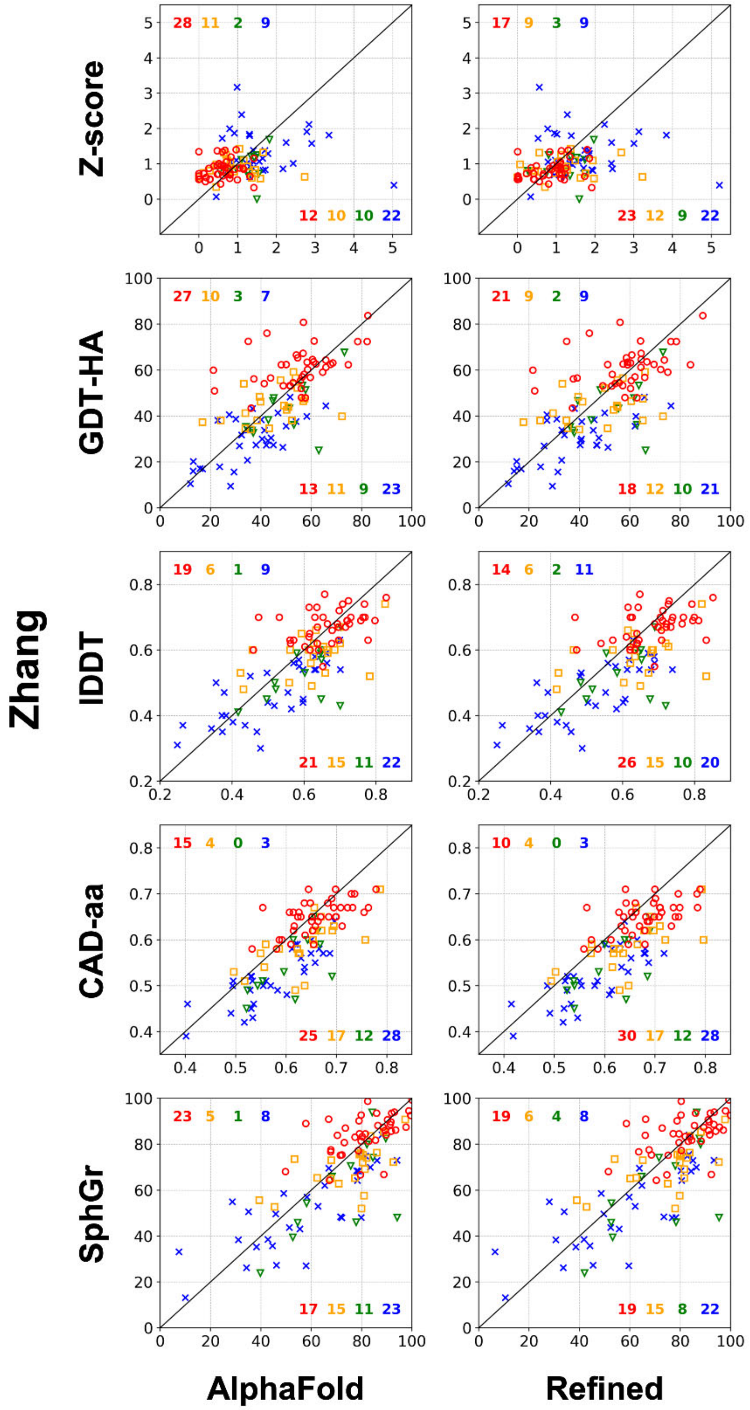


**Figure 1 – Supplementary 3.** Head-to-head comparison between initial and refined AlphaFold models and predictions from Zhang, the next best prediction group in CASP!3, in terms of different quality measures. *See* **Figure 1B** and **Figure 1 - Supplementary 1**.

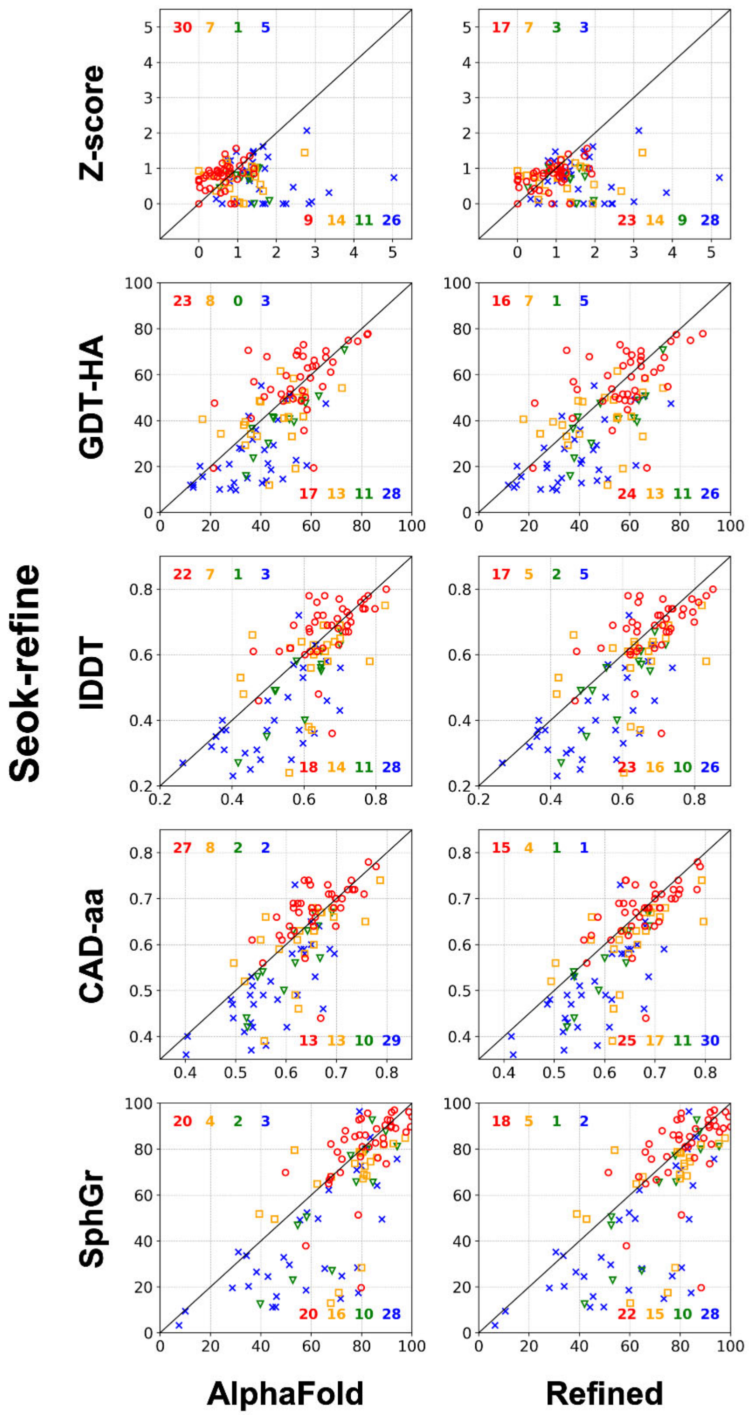


**Figure 1 – Supplementary 4.** Head-to-head comparison between initial and refined AlphaFold models and models generated by Seok-refine during CASP13 in terms of different quality measures. *See* **Figure 1B** and **Figure 1 - Supplementary 1**.

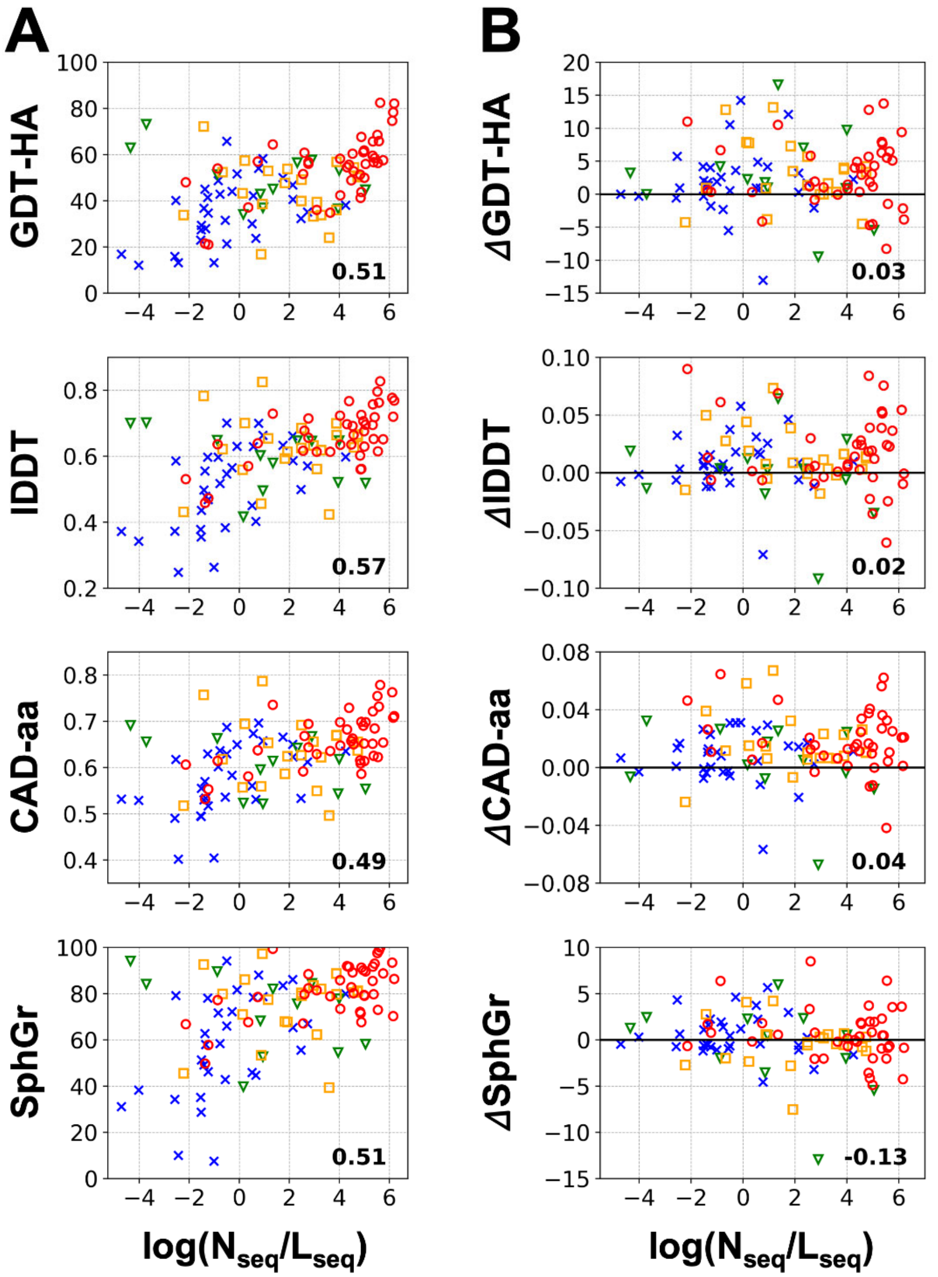


**Figure 1 – Supplementary 5.** (A) AlphaFold prediction accuracy vs. the number of homologous sequences available for a given target. (B) Changes in quality measures after refinement as a function of homologous sequences. Symbols and colors distinguish different target categories as in **Figure 1B**. Pearson's correlation coefficients from linear regression fits are indicated for each plot.
